# Decoupling gene knockout effects from gene functions by evolutionary analyses

**DOI:** 10.1101/688358

**Authors:** Li Liu, Mengdi Liu, Di Zhang, Shanjun Deng, Piaopiao Chen, Jing Yang, Yunhan Xie, Xionglei He

**Affiliations:** State Key Laboratory of Biocontrol, School of Life Sciences, Sun Yat-sen University, Guangzhou 510275, China

## Abstract

Genic functions have long been confounded by pleiotropic mutational effects. To understand such genetic effects, we examine HAP4, a well-studied transcription factor in *Saccharomyces cerevisiae* that functions by forming a tetramer with HAP2, HAP3, and HAP5. Deletion of HAP4 results in highly pleiotropic gene expression responses, some of which are clustered in related cellular processes (clustered effects) while most are distributed randomly across diverse cellular processes (distributed effects). Strikingly, the distributed effects that account for much of HAP4 pleiotropy tend to be non-heritable in a population, suggesting they have little evolutionary consequences. Indeed, these effects are poorly conserved in closely related yeasts. We further show substantial overlaps of clustered effects, but not distributed effects, among the four genes encoding the HAP2/3/4/5 tetramer. This pattern holds for other biochemically characterized yeast protein complexes or metabolic pathways. Examination of a set of cell morphological traits of the deletion lines yields consistent results. Hence, only some gene deletion effects support related biochemical understandings with the rest being pleiotropic and evolutionarily decoupled from the gene’s normal functions.

## Introduction

Mutation analysis has long been used to understand the functions of a gene(*1*). It appears now clear that a gene can often affect various seemingly unrelated traits(*2*), a phenomenon termed pleiotropy(*3*). For instance, a large-scale gene knockdown assay in the nematode worm *Caenorhabditis elegans* finds on average a gene affects ∼10% of 44 assessed traits(*4*). Attempts to understand such pleiotropic mutational effects are mainly from mechanistic perspectives(*5*, *6*), by considering the focal gene’s multiple molecular functions or multiple cellular processes associated with a single molecular function(*7*). The resulting pictures are, however, often complex, confusing our understandings in how a gene functions.

Since biological systems are all evolutionary products with history, mechanistic perspectives alone may bias the efforts for delineating a biological phenomenon(*8*, *9*). This is exemplified by the debates on the ENCODE project in which up to 80% of the human genome was claimed to be functional despite that only 10% appears to be under selection(*10*–*12*). A simple example explains how the confusion arose. Suppose there is a transcription factor (TF) that recognizes a DNA motif, say, ATCGATC. The human genome with ∼3×10^9^ base pairs in length contains over one hundred thousand ATCGATC motif sequences, some of which are evolutionarily selected for certain biological processes while the rest exist as *ad hoc* entities due to the equilibrium of random mutations in such a long genome(*11*). From a purely mechanistic perspective originally adopted by the ENCODE consortium(*10*), the myriad ATCGATC sequences are all called functional so long as they are bound by the TF. However, the claim of *ad hoc* entities as functional would only confuse our understandings in how the system is organized to function. Such confusions forced the ENCODE consortium to eventually abandon their evolution-free view on biochemical functionality(*13*). Notably, the same problem actually also applies to the genetic effects defined in reverse genetics analysis. The common practice is to knockout or knockdown a gene and find the traits significantly altered(*14*), which represents a purely mechanistic framework. In the above TF versus ATCGATC motif example, when the TF is deleted, the expression of those genes with the motif at promoter region could all be affected. The resulting pleiotropic effects, which are either *ad hoc* or evolutionarily selected according to the nature of the focal motifs, together would lead to a very complex picture on the functionality of the TF.

The necessity of adopting an evolutionary view in reverse genetic analysis lies also in the effect size of the knockout or knockdown mutations experimentally introduced, which is much larger than that of typical segregating alleles in natural populations(*15*). Hence, while the normal functions of a gene is necessarily built by natural selection, responses to such experimental inactivation of the functions may not be shaped by evolution(*16*). Then, how can the “non-evolutionary” responses be in line with the evolutionarily selected gene functions? With this question in mind we here examine the evolutionary nature of a set of gene deletion effects. We show widespread decoupling of gene deletion effects from the gene’s normal functions, calling for an evolutionary framework for reverse genetic analysis.

## Results

We started with a known yeast *Saccharomyces cerevisiae* gene HAP4(*17*). It is a non-essential transcription factor that has been subjected to extensive studies since its discovery 30 years ago(*18*). We deleted the open reading frame of HAP4 in *S. cerevisiae* strain BY4741, and checked the expression trait of the other yeast genes by sequencing the transcriptome of the strains that grow in the rich medium YPD at 30 °C (Table S1). We found 195 responsive genes each with a significant expression change under a stringent statistical cutoff (Table S2). Gene ontology (GO) analysis of the 195 genes revealed one third of them (65/195) clustered more than expected by chance in dozens of GO terms. These GO terms are related with each other and reflect well the functional annotations of HAP4 as a regulator of mitochondria activities(*19*) (Fig. 1A). The remaining two thirds (130/195) distribute rather randomly across diverse biological processes, underscoring the strong pleiotropy of HAP4. The two sets of genes are all functionally characterized with clear GO annotations (Table S2), and have comparable fitness importance (*P* = 0.1, Mann-Whitney U-test; Fig. S1). Notably, the 65 clustered deletion effects and 130 distributed deletion effects are supported by similar P-values and fold changes (*P* = 0.20 and 0.46, respectively, Mann-Whitney U-test; Fig. 1B).

**Fig. 1.**
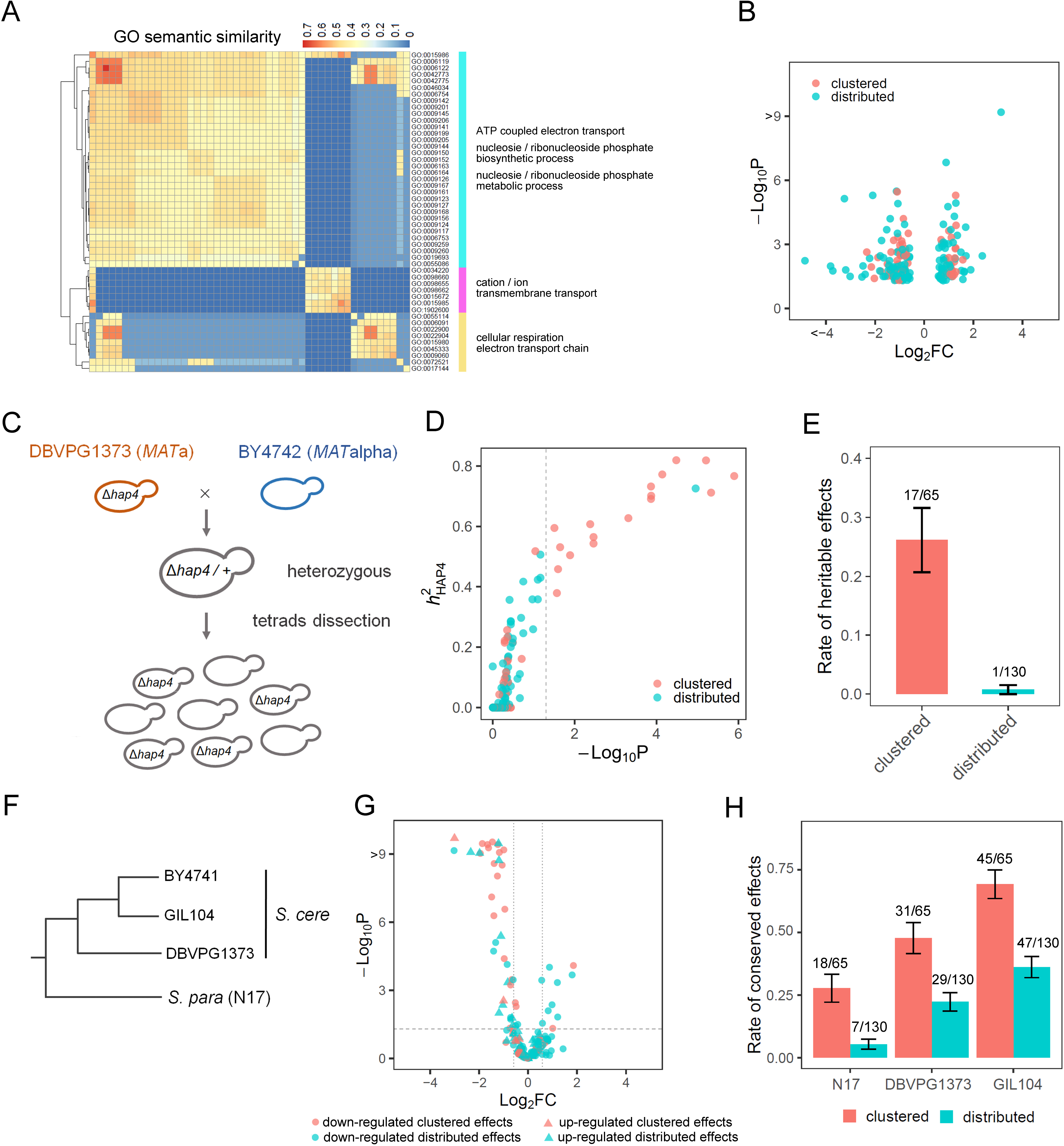
Tests of evolutionary effectiveness of HAP4 deletion effects. **(A):** The 195 responsive genes are enriched in dozens of GO terms that are related and also reflect well the functional annotations of HAP4. The heatmap shows the pairwise similarity of the enriched GO terms, with three subclasses each corresponding to certain biological processes that are summarized at the right. **(B):** The *P*-value (adjusted for multiple testing) and fold change (FC) of the 65 clustered effects and 130 distributed effects. Each dot represents a responsive gene (i.e., an effect). **(C):** Obtain a population of segregants with different genetic backgrounds to test the heritability of HAP4 deletion effects. **(D):** The 65 clustered effects have greater *h*^2^_HAP4_ than the 130 distributed effects (*P* = 7.8×10^−4^, Mann-Whitney U-test). Each dot represents an effect, and *P*-value measures the statistical significance of *h*^2^_HAP4_, with the vertical dashed line showing adjusted *P* = 0.05. **(E):** The proportions of deletion effects that are significantly heritable with adjusted *P* < 0.05. Error bars represent SE. **(F):** A dendrogram showing the phylogeny the four yeast strains examined in this study. **(G):** Conservation analysis of the HAP4 deletion effects. The 195 responsive genes defined in BY4741(*Δhap4*) are examined with respect to their expression responses in *S. paradoxus* N17 (*Δhap4*). The horizontal dashed line shows adjusted *P* = 0.05 and vertical dashed lines show log_2_FC = ±0.58. (cyan: clustered effects; red: distributed effects; cycle: down-regulated in BY4741(*Δhap4*); triangle: up-regulated in BY4741(*Δhap4*)) **(H):** The rate of conservation in the three related yeasts for the 65 clustered effects and 130 distributed effects defined in BY4741(*Δhap4*), respectively. Error bars represent SE.

Because evolution happens in a population rather than in an individual, it is important to test the population-level heritability of the deletion effects. We crossed two *S. cerevisiae* strains to obtain a population of yeast segregants. Specifically, a wild-type strain BY4742(*MATalpha*), which is identical to BY4741 except at the mating locus, was crossed with the HAP4 deletion line of DBVPG1373(*MATa*) (Fig. 1C). This way, the comparison between wild-type and null alleles at the HAP4 locus would match the comparison conducted in the isogenic BY4741 background. We dissected six tetrads of the hybrid and obtained 12 HAP4 wild-type and 12 HAP4 null segregants. For each of the 195 deletion effects we computed its heritability (*h*^2^_HAP4_) in the segregant population (Methods). The *h*^2^_HAP4_ measures the fraction of variance of an expression trait that is attributed to the HAP4 locus. We noted that the heritability analysis resembles a forward genetic assay with a candidate genetic locus. An *h*^2^_HAP4_ close to zero suggests HAP4 is not a QTL (Quantitative Trait Locus) of the expression trait. We found the 65 clustered effects in general have much higher *h*^2^_HAP4_ than the 130 distributed effects (*P* = 7.8×10^−4^, Mann-Whitney U-test; Fig. 1D). Approximately 26.2% (17/65) of the clustered effects have a statistically significant *h*^2^_HAP4_, while the number is 0.8% (1/130) for the distributed effects (*P* = 2.1×10^−8^, Fisher’s exact test; Fig. 1E). It is worth pointing out that mutational effects sensitive to genetic backgrounds have been documented in a wide range of organisms(*20*–*23*).

Because non-heritable phenomena in biology last only one generation, with little evolutionary consequences, the deletion effects with low population-level heritability should be evolutionarily unconserved. We obtained, under the same environment and the same statistical cutoff as in BY4741, the deletion effects of HAP4 in *Saccharomyces paradoxus* strain N17, a closely related yeast species diverged from *S. cerevisiae* ∼10 million years ago(*24*) (Fig. 1F). Only 5.4% (7/130) of the distributed effects found in *S. cerevisiae* were also observed in the HAP4 deletion line of *S. paradoxus* N17, while the number was 27.7% (18/65) for the clustered effects (*P* = 3.1×10^−5^, Chi-square test). The difference was robust as evidenced by plotting the expression responses of the 195 genes in N17(*Δhap4*) (Fig. 1G). Notably, although the statistical signals were not directly comparable between the conservation analysis and heritability analysis, there were 14 overlaps between the 25 conserved effects defined here and the 18 effects with significant *h*^2^_HAP4_. Hence, the heritability analysis based on a rather arbitrary and small population appeared to represent well the situation in nature. We also examined intra-species conservation of the HAP4 deletion effects by looking at two other *S. cerevisiae* strains DBVPG1373 and GIL104, the former of which is 0.35% diverged from the strain BY4741 and the latter 0.07% diverged at the genomic level(*25*). Similar1y, the 130 distributed effects were largely unreproducible in strains DBVPG1373(*Δhap4*) and GIL104(*Δhap4*), while the 65 clustered effects were much more conserved (*P* = 0.5×10^−4^ for DBVPG1373(*Δhap4*) and *P* = 2.6×10^−5^ for GIL104(*Δhap4*), Chi-square test; Fig. S2). As expected, both types of the deletion effects are more reproducible in more related yeasts (Fig. 1H). We confirmed the results cannot be explained by different detectability of expression changes between the two gene sets by excluding those genes lowly expressed in wild-type BY4741 (Fig. S3). These data, together with the heritability analysis, suggest the clustered effects of HAP4 tend to be evolutionarily selected; on the contrary, the distributed genetic effects appear largely non-evolutionary, likely representing *ad hoc* responses to the gene deletion(*16*).

HAP4 functions by forming a tetramer with HAP2, HAP3 and HAP5, which is a result of evolution(*18*). We hypothesized clustered effects should support this biochemistry understanding better than distributed effects, because the latter is non-evolutionary. To test the hypothesis, we deleted the other three genes that encode the tetramer, respectively, in *S. cerevisiae* BY4741, and measured the expression profiles of the deletion lines. We defined clustered effects and distributed effects for each of the lines using the same method as in the HAP4 deletion line. We obtained 43, 150 and 50 clustered effects, and 61, 306 and 111 distributed effects for the deletions of HAP2, HAP3 and HAP5, respectively (Fig. 2A-C; Table S3). Consistent with the hypothesis, we found 20 overlapped clustered effects across the four gene deletion lines, 14.5 times higher than that of the distributed effects (*P* < 0.001, simulation test, Fig. 2D). Notably, the 20 overlapped clustered effects are not the strongest in BY4741(*Δhap4*) (Fig. S4). To avoid the potential bias that expression responses to the tetramer may have been considered in the GO annotations of the responsive genes, we excluded all expression-related evidences for GO annotations to define new clustered and distributed effects. We obtained essentially the same result (Fig. S5). Because there are publicly available microarray data for HAP2, HAP3, HAP4 and HAP5 deletion lines(*26*), we also repeated the analysis using the public expression data and observed a similar pattern (Fig. S6).

**Fig. 2.**
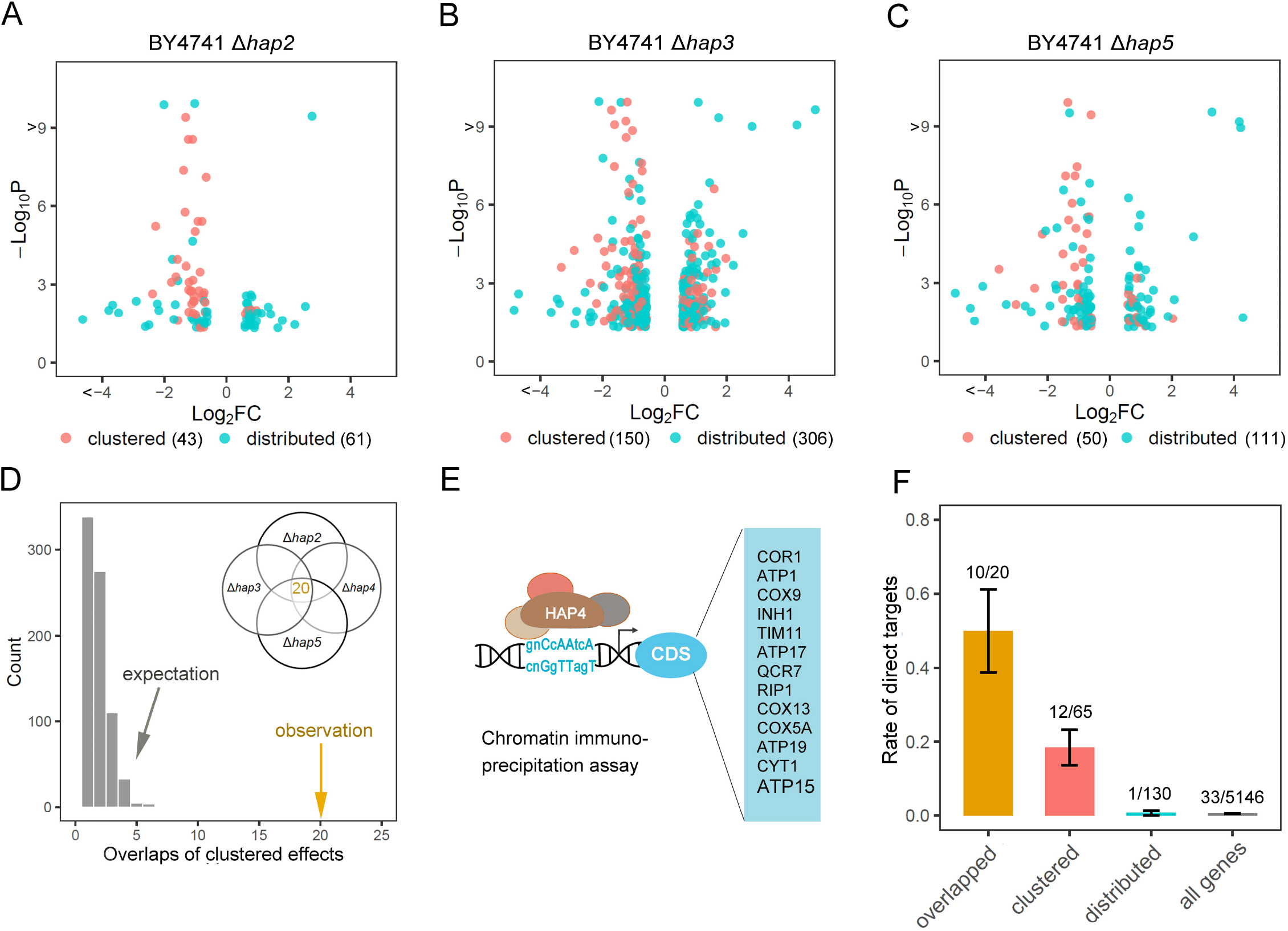
Clustered effects of the four genes encoding the HAP2/3/4/5 tetramer overlap a lot more than their distributed effects do. **(A, B, C):** The P-value (adjusted for multiple testing) and fold change (FC) of the clustered effects and distributed effects defined in the three BY4741 strains *Δhap2*, *Δhap3*, and *Δhap5*, respectively. Each dot represents a responsive gene (i.e., an effect), and the total number of responsive genes is shown at the bottom next to the effect type. **(D):** There are 20 overlapped clustered effects for the four genes encoding the HAP2/3/4/5 tetramer, which is significantly higher than expectation. The expectation is estimated by random sampling of the distributed effects of the four genes to calculate overlaps, and 1,000 such simulations were conducted. **(E):** Among the 195 responsive genes found in BY4741(*Δhap4*) 13 are direct target of HAP4 according to a chromatin immune-precipitation assay. **(F):** The proportion of direct target of HAP4 in different gene sets. Error bars represent SE.

In addition to considering the protein complex formed by HAP4, we could also consider protein-DNA interactions since HAP4 is a transcription factor. Data from a chromatin immuno-precipitation (CHIP) assay of the promoters bound by HAP4 show that, among the 195 responsive genes observed in BY4741(*Δhap4*), 13 are direct targets of HAP4 (Fig. 2E)(*27*, *28*). Interestingly, there is 24-fold enrichment of direct targets in the clustered effects relative to the distributed effects (*P* = 8.6×10^−6^, Fisher’s exact test); among the 20 overlapped clustered effects 50% (10/20) are direct targets of HAP4, while the genomic background is 0.64% (33/5146) (*P* = 4.6×10^−18^, Hypergeometric test) (Fig. 2F). Hence, the CHIP data well support the distinction of the two effect types.

Collectively, these results are consistent with a previous model(*16*) (Box 1), in which the null phenotype of a gene can be ascribed to either the loss of the gene’s native functions, or the gain of spurious functions that arise from passive adjustments of the cellular system after the perturbation. The key difference of the two function types is their evolutionary nature: native functions are historical, selected, and evolutionary, while spurious functions are ahistorical, *ad hoc*, and non-evolutionary (*29*–*31*). Accordingly, the distributed effects examined here likely represent spurious functions created by the HAP4 deletion, and the clustered effects could be in a large part ascribed to the native functions of HAP4. It is intriguing how the two effect types defined by GO could fit the two function types described in the model. We reasoned that evolutionarily optimized native functions are likely to regulate specific pathways or processes; losing them would thus cause coordinating changes of the related genes(*9*), which are detected by GO analysis. In contrast, spurious functions may affect the transcriptome in a rather random way, resulting in distributed changes across diverse cellular processes, most of which cannot be covered by overrepresented GO terms. This may explain why GO clustering here could echo evolutionary effectiveness.

### Box 1

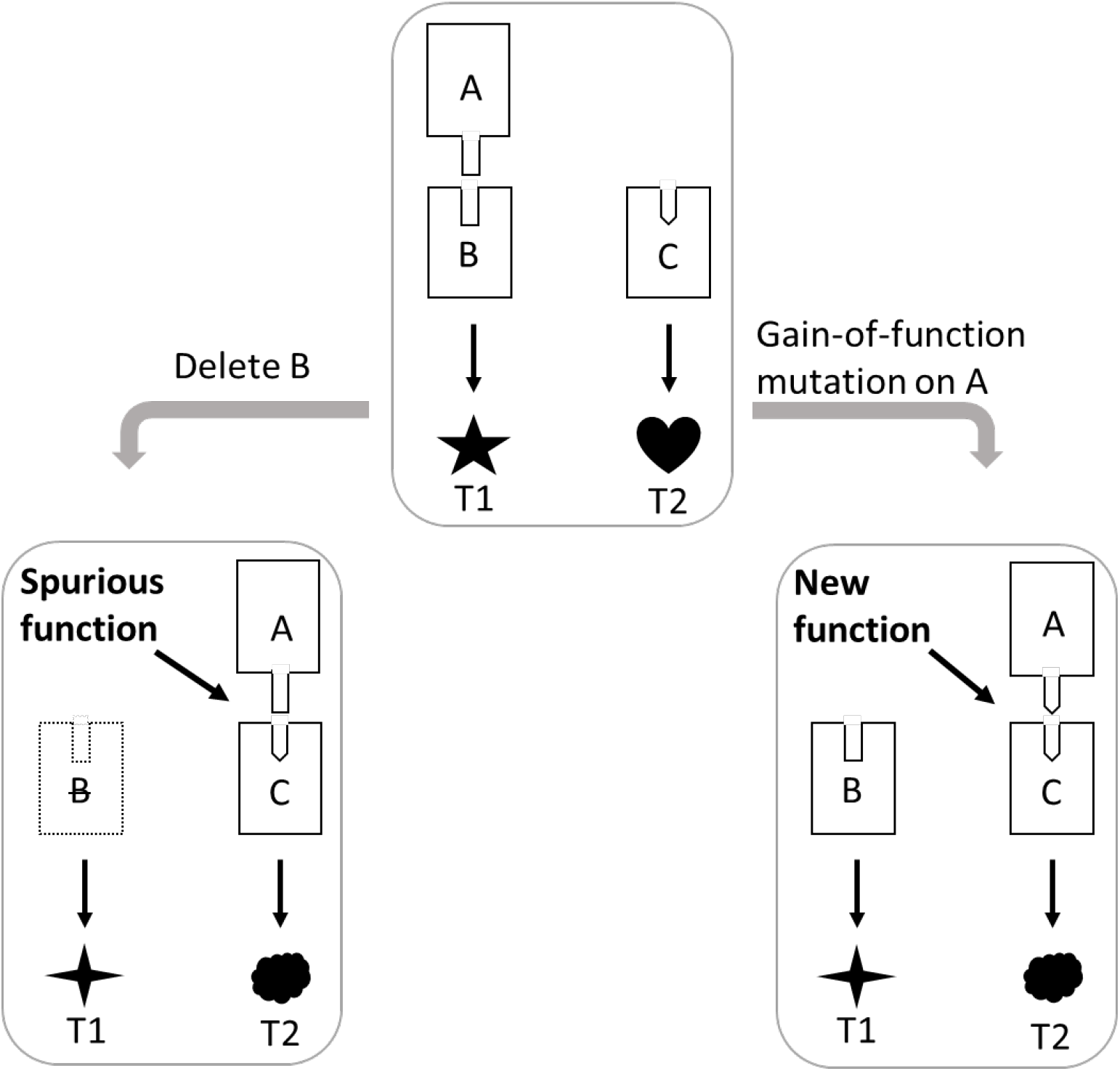

Nearly 90 years ago Hermann J. Muller coined the terms amorph, hypermorphy, hypomorph, antimorph, and neomorph to classify mutations based on their loss- or gain-of-function nature (see ref. 1). The basic idea of the classification has been fundamental to genetic analysis. In particular, **amorph** refers to null mutation on a gene, and the resulting phenotype is believed to represent the gene’s native function. In contrast, **neomorph** refers to gain-of-function mutation on a gene, and the resulting phenotype does not represent the gene’s native function. These concepts, although intuitively valid, lack rigorous tests. The diagram on the left illustrates how confusion could arise.

Suppose there is a living system with three genes (A, B and C) and two traits (T1 and T2). The function of proteins A and B is to form a dimer to regulate T1, and the function of protein C is to regulate T2. The understand the system we may apply genetic analysis. Deletion of B will break the A-B dimer, altering T1. This phenotype change represents the **native function** of B. However, when B is absent, A may find C to form a new, although less intimate, dimer A-C, which would alter T2. This is plausible since proteins with a structurally similar domain are prevalent in a eukaryotic genome. The change of T2 does not represent the native function of B; instead, it is explained by the non-native A-C dimer, a **spurious function** that arises from the deletion of B.

Notably, the spurious function arising from the deletion of B is by nature same as the **new function** caused by a gain-of-function mutation on A. It is well accepted that phenotype changes resulting from gain-of-function mutations do not represent native functions. From an evolutionary perspective only the A-B dimer is **selected**; the A-C dimer in both mutated systems is ***ad hoc***.

Regardless of the underlying logic, clustered genetic effects seem to be well matched with related biochemical understandings. This would help address a long-standing challenge to molecular biology - the gap between genetic analysis and biochemistry analysis(*14*, *32*); specifically, genes with intimate biochemical interactions do not have common genetic effects and genes with common genetic effects do not show intimate biochemical interactions(*33*, *34*). To test the generality of the finding that was based on the HAP2/3/4/5 tetramer, we examined other biochemically characterized protein complexes by using publicly available expression data. To avoid bias we considered a single dataset comprising microarray-based expression profiles of over one thousand yeast gene deletion lines(*26*). There are 54 protein complexes annotated by a previous study suitable for our analysis(*35*). In 24 cases the overlaps of clustered effects are significantly more than what would be expected from distributed effects at a 99% confidence level, and the enrichments range from 2.7-fold to over 100-fold with a median 5.3-fold (Fig. 3A and Table S4). The overlapped clustered effects of each protein complex represent specific functions (Fig. 3B-C and Fig. S7). For example, the ten overlapped clustered effects of the elongator holoenzyme complex are tens to hundreds times overrepresented in a few transcription-related GO terms as well as proteasome-related GO terms (Fig. 3B), the former of which echo well the annotated functions of the complex while the latter appear to suggest new understandings(*36*). As another example, the genes encoding the protein kinase CK2 complex have 21 overlapped clustered effects that appear to affect specifically the metabolism of various amino acids (Fig. 3C), a functional insight not been well recognized(*37*).

**Fig. 3.**
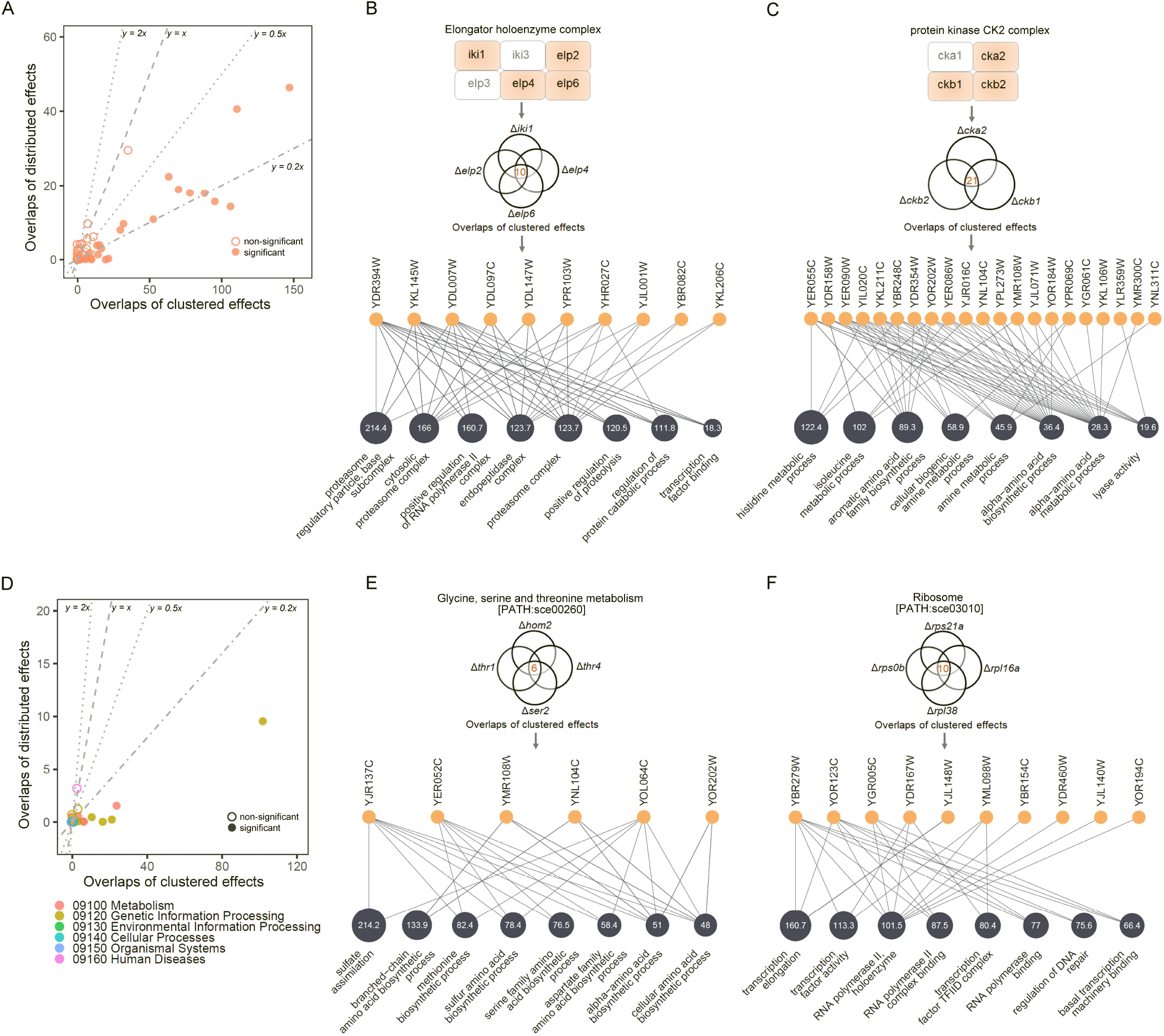
Clustered effects support related biochemistry understandings much better than distributed effects in a variety of protein complexes and KEGG pathways. **(A):** The clustered effects of genes encoding a protein complex in general overlap more than their distributed effects. Each circle represents a complex, and the filled ones are significant at a 99% confidence level estimated by random sampling. A total of 54 protein complexes are included here, with 24 cases showing at least twice more overlapped clustered effects than overlapped distributed effects (below the line y = 0.5x). The numbers of effects have been normalized such that in each case the overlaps of clustered effects and the overlaps of distributed effects can be directly compared. **(B):** The representative GO terms of the overlapped clustered effects of the elongator holoenzyme complex. Only four genes encoding the complex, which are highlighted in orange, have suitable expression data for the analysis. There are 10 overlapped clustered effects of the four genes, which are over 300 times more than expected. The expectation is estimated by random sampling of the distributed effects of the focal genes to calculate overlaps. The orange circle each represents an overlapped clustered effect, and the blue circles represent the enriched GO terms of the overlapped clustered effects with the number inside showing the fold enrichment in the given term. **(C):** The representative GO terms of the overlapped clustered effects of the protein kinase CK2 complex. There are 21 overlapped clustered effects, 83.3 times more than expected. **(D):** The clustered effects of genes in the same KEGG pathway also tend to overlap more than their distributed effects. Each circle represents a pathway, and the filled ones are significant at a 99% confidence level. A total of 41 pathways are included, with nine showing at least five times more overlapped clustered effects than overlapped distributed effects (below the line y = 0.2x). The number of effects have been normalized such that in each case the overlaps of clustered effects and the overlaps of distributed effects can be directly compared. **(E):** The representative GO terms of the overlapped clustered effects of the four genes in the metabolic pathway sce00260. There are six overlapped clustered effects, over 100 times more than expected. **(F):** The representative GO terms of the overlapped clustered effects of the four genes in the genetic information processing pathway sce03010. There are 10 overlapped clustered effects, 21.4 times more than expected.

We also checked genes on the same KEGG pathways. There are 41 pathways that are related to metabolism, genetic information processing, cellular processes, and so on, suitable for our analysis (Table S5). The rate of overlaps of clustered effects is significant higher than that of distributed effects in nine cases, and the enrichments range from 5.6-fold to over 100-fold with a median 46.9-fold (Fig. 3D and Table S5). Consistently, the overlapped clustered effects of each pathway represent distinct functions (Fig. 3E-F and Fig. S7). For the many cases in which clustered effects show no more overlaps than distributed effects, the involved genes may execute additional functions irrelevant to the focal complex or pathway. Notably, in none of the cases distributed effects represent related biochemical understandings better than clustered effects, highlighting the cryptic nature of them. Taken together, focusing on clustered effects appears to be a readily operational approach to narrowing the gap between genetic analysis and biochemical data.

The above analyses considered gene expression traits. We next examined the yeast cell morphological traits that are based on the microscopic images of cells stained by fluorescent dyes(*38*). With the help of a computer software as many as 405 quantitative traits can be obtained from cell wall and nuclear stained cell images(*39*). These traits are typically about area, distance, and angle calculated based on dozens of coordinate points, lines and angles that describe the shape of mother cell and bud, and the shape and localization of the nuclei in mother cell and bud (Fig. 4A). This large set of yeast traits had served as a valuable resource for studying genotype-phenotype relationships(*9*, *40*, *41*). Deletion of HAP4 in *S. cerevisiae* significantly altered 78 morphological traits, among which 24 are also significantly affected in *S. paradoxus* by HAP4 deletion (Table S6). To test if the evolutionarily conserved effects of HAP4 are shared with HAP2, HAP3 and HAP5 more than the non-conserved effects, we also measured the morphological traits affected by each of the other three genes, respectively, in *S. cerevisiae*. We found that 58.3% (14/24) of the conserved effects are shared with all the other three genes, which is significantly higher than the number (18/54 = 33.3%) for the non-conserved effects of HAP4 (*P* = 0.035, Fisher’s exact test; Fig. 4B). The estimations are not explained by correlated traits (Fig. S8), and the difference remains largely unchanged when only traits with small measuring noise are considered (Fig. S9). Hence, the cell morphology data also support the role of evolution in separating genetic effects.

**Fig. 4.**
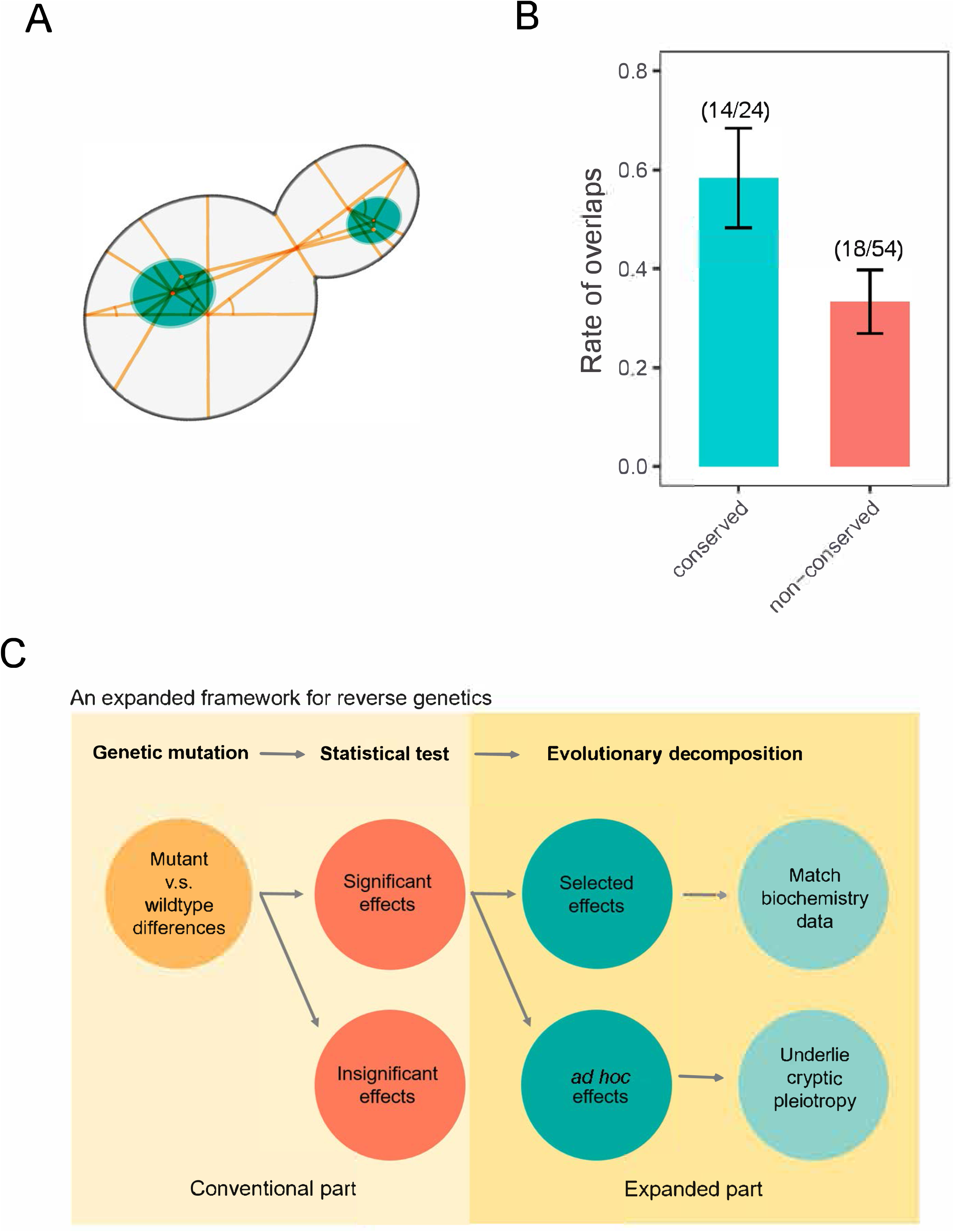
Examination of cell morphological traits also supports the role of evolution in separating genetic effects. **(A):** The yeast cell morphology outlined by coordinate points, lines and angles (only some are shown) based on which a total of 405 quantitative traits can be derived by a computer software. **(B):** The traits affected by HAP4 in both *S. cerevisiae* and *S. paradoxus* (i.e., conserved effects) are more likely to overlap with those affected by HAP2, HAP3 and HAP5 than the traits affected by HAP4 only in *S. cerevisiae* (non-conserved effects) (*P* = 0.035, one-tailed Fisher’s exact test). A total of 78 morphological traits significantly affected by HAP4 deletion in *S. cerevisiae* are examined, among which 24 are conserved effects and 54 non-conserved effects. Overlaps refer to traits significantly affected by all four gene deletions in *S. cerevisiae.* Error bars represent SE. **(C):** Proposition of an expanded framework for reverse genetic analysis. Statistically significant genetic effects defined in conventional framework are further separated into evolutionarily selected and *ad hoc* ones, with the former supporting related biochemistry understandings and the latter being pleiotropic and decoupled from the gene’s normal functions.

## Discussion

Thanks to the mature framework of measuring the selective constraints on DNA sequence(*42*), the evolution-free functionality of DNA elements defined in ENCODE was challenged immediately after its emergence(*11*, *12*). Notably, the gene-trait interactions defined in reverse genetic analyses are also based on an evolution-free framework. However, this century-old problem has been largely ignored, despite exceptions(*43*, *44*), primarily due to the lack of a readily available measure of the underlying evolutionary constraint. In this study we performed, for the first time to the best of our knowledge, a rigorous test of the evolutionary nature of a set of gene deletion effects by examining their within-population heritability and intra-/inter-species conservation. We found only some of them subject to effective selection, with the rest likely being *ad hoc* and non-evolutionary. That being said, we cautioned some effects might be under very weak selection that was beyond the detection power of our analyses. This concern would be alleviated by a reasonable assumption that effects under very weak selection are not distinct from those under no selection in the functional properties examined in the study. Similar to the *ad hoc* “functional” DNA elements defined in ENCODE(*10*), the *ad hoc* genetic effects are presumably explained by mutation equilibrium or spurious functions arising from the gene deletion(*16*). Importantly, since such *ad hoc* effects have not yet been shaped by evolution, they are unlikely to be compatible with the roles the focal gene has long played in evolution(*9*). This may explain in great part the origin of gene pleiotropy.

Conceptually speaking, our evolutionary view on genetic effects is an extension of the evolutionary view on the biochemical activities of DNA elements(*11*, *12*). Hence, pros and cons that have widely discussed in the debates on the ENCODE project apply similarly to this study. For example, because detecting selection involves multiple lineages, one cannot rule out the possible that an entity under no detectable selection is actually subject to lineage-specific selection(*11*). However, since the lineages examined are often closely related, lineage-specific entities selected in a short time window should be rare compared to those acquired during the long time period predating the split of the lineages. Operationally speaking, the evolutionary view on the functionality of DNA elements relies on DNA sequence comparison, which is straightforward and now mature. However, an evolutionary separation of genetic effects requires rather complex experimental designs; also, there is no available framework for modeling the turnover rate of gene-trait interactions under no selection. Hence, we could, as in this study, only perform enrichment analysis for a group of genetic effects. Nevertheless, the current limitation in operationality does not challenge the validity of the concept. A surprising finding of this study is GO clustering can serve as a useful and readily operational proxy for selection when expression traits are examined. The underlying rationale, namely, functional coordination built by selection, may help us design more efficient strategies for delineating the evolutionary nature of genetic effects in the future.

In summary, by examining the evolutionary nature of a set of gene deletion effects we revealed widespread decoupling of gene deletion effects from gene functions. This calls for an expanded framework for reverse genetic analysis (Fig. 4C). Specifically, the conventional framework relies solely on statistical tests to separate the mutant versus wild-type differences into significant and insignificant effects. In the expanded framework significant effects are further separated into evolutionarily selected and evolutionarily *ad hoc* ones. Only the former would support the biochemical understandings with the latter being pleiotropic and decoupled from the gene functions.

## Materials and Methods

### Yeast gene deletions

Three *S. cerevisiae* (SC) strains BY4741 (*MATa, his3, leu2, met15, ura3*), GIL104 (*MATa, URA3, leu2, trp1, CAN1, ade2, his3, bar1Δ::ADE2*; derived from the W303) and DBVPG1373 (*MATa, ura3*), and one *S. paradoxus* (SP) strain N17 (*MATa, ura3*) were included in the study. Unless otherwise stated, the *S. cerevisiae* strains were cultured in the rich medium YPD (1%Yeast extract, 1%Peptone, 2% Dextrose) at 30°C, and *S. paradoxus* N17 was cultured in YPD at 25°C. The wild type URA3 in GIL104 was first replaced by a LEU2 cassette. HAP4 was replaced by a URA3 cassette in each of the four strains. HAP2, HAP3, and HAP5 were also replaced, respectively, by a URA3 cassette in BY4741. Notably, for all gene replacements the whole open reading frame from the starting codon to the stop codon of a focal gene was replaced. As described in our previous study(*22*), the standard LiAc transformation method(*45*) was used to transform DNA into the yeast cells and gene replacements were achieved by homologous recombination. The transformation protocol was slightly modified for *S. paradoxus*(*46*); specifically, heat shock was performed for seven minutes at 37°C. Synthetic medium deprived of uracil or leucine was used to select the clones with successful replacement for the target gene. All gene replacements were verified using polymerase chain reaction (PCR). For each gene deletion line, 3-5 independent clones were obtained for further examination, which effectively controlled the potential effects of secondary mutations introduced during the gene replacement.

Because haploid yeast cells tend to flocculate, which is not suitable for cell morphology characterization, diploid yeasts are required in the analysis of morphological traits. We first obtained haploid gene deletion strains (SC-BY4741 or SP-N17 background; *MATa*), which were then crossed with the corresponding *MATalpha* wild-type strain, respectively. The diploid heterozygous gene deletion strains were sporulated by following the method of a previous study(*47*). Specifically, the cells were incubated in YEP (1% yeast extract,1% Bacto peptone,0.05% NaCl) containing 2% potassium acetate) for five hours at 30 °C to start the sporulation process. The culture was centrifuged (2,000g, for 2 min), the cell pellet was washed three times by sterile water, and re-suspended in sporulation media (10g/l potassium acetate and 50mg/l zinc acetate) for five days at 25°C with shaking. The products were incubated with 200U/ml lyticase (Sigma #L4025) for 30 min at 30°C followed by 15 min at 50°C. The products were washed by sterile water and then plated on synthetic medium deprived of uracil for two days at 30°C for SC strains, or 25°C for SP strains. The genotypes of the colonies were determined by PCR. For each gene the haploid deletion strains of both mating types were obtained. A pair of *MATa* and *MATalpha* strains with the same gene deletion were crossed to obtain a diploid homozygous gene deletion strain. For each gene deletion line three independent clones were obtained and examined.

### Obtain a population of segregants

A wild-type strain BY4742 (*MATalpha*), which is identical to BY4741 except at the mating locus, was crossed with a HAP4 deletion stain of DBVPG1373 (*MATa*). Two biological replications were carried out. The diploid heterozygous deletion strains were sporulated for 3-5 days in sporulation medium on a shaking table at 25°C. Tetrads were obtained and incubated with 200U/ml lyticase for 3-5 min at 30°C, and then streaked onto a YPD plate for tetrad dissection using the MSM400 dissection microscope (Singer Instrument Company Ltd). Spores were grown on YPD plates at 30°C for two days, and the genotypes of the colonies were determined by PCR. We selected only those tetrads that produce four segregants with genotypes *MATa*+HAP4, *MATa*+*Δhap*4, *MATalpha*+HAP4, and *MATalpha*+*Δhap4*, respectively. A total of 24 segregants from six such tetrads were obtained for the heritability analysis.

### RNA sequencing and data analysis

For each strain a single colony on agar plate was picked and cultured in YPD liquid overnight at 30 °C with shaking. Approximately 200μl saturated culture was added into 10ml fresh YPD, which resulted in an optical density OD600 ∼ 0.1 (UNICO UV/VIS Spectrophotometer), and cells of 3ml culture at OD600 = 0.5-0.65 were harvested. Total RNA was extracted by QIAGEN RNeasy Plus mini kit (Cat No.74136). The mRNA sequencing was performed using the paired-end module on a HiSeq platform at Genwiz by following the standard procedure. To ensure the high quality of expression analysis, we sequenced the mRNA of 3-6 independent clones for each wild-type or gene deletion line.

RNA-seq reads were mapped to reference yeast genomes using STAR (Version 2.6.0c)(*48*). For BY4741 and GIL104, we used the genome of *S. cerevisiae* strain S288C as the reference (version R64-2-1_20150113; http://www.yeastgenome.org). The reference genomes of SC-DBVPG1373 and SP-N17 were downloaded from SGRP (https://www.sanger.ac.uk/research/projects/genomeinformatics/sgrp.html). For a typical clone there were about 6.5 million paired-end reads mapped to the coding sequences. Gene expression levels were determined by Featurecounts (version 1.6.2)(*49*) with default settings and RPKM (reads per kilobase per million) of each gene were calculated by R package edgeR. The wild-type versus mutant differential expression analysis was performed by DESeq2(*50*) with default parameters, and genes with an adjusted P-value smaller than 0.05 and a fold change (FC) greater than 1.5 were defined as significantly changed genes. In the conservation, heritability or overlapping analysis, an effect is called conserved, heritable or overlapped only when it shows the same direction in the various conditions examined. Genes of the uracil biosynthesis pathway (YBL039C, YEL021W, YJL130C, YJR103W, YKL024C, YKL216W, YLR420W, YML106W, YMR271C) were excluded from further analysis. The expression level measured by RPKM of HAP2, HAP3, HAP4 and HAP5 in wild-type BY4741 is 84.4, 64, 194 and 90.5, respectively, and all becomes zero in the corresponding deletion lines. The fitness of yeast gene deletion lines was produced by a previous study(*51*). Table S1 contains details of the RNA-seq expression information of all yeast lines examined in this study.

### GO analysis

The GO analyses of the responsive genes derived from our RNA-seq data were conducted in the SGD website using GO Term Finder (Version 0.86; https://www.yeastgenome.org/goTermFinder), by excluding computational analysis evidences and other less reliable evidences: IBA, IC, IEA, IKR, IRD, ISA, ISM, ISO, ISS, NAS, ND, TAS. In a strict analysis which required the exclusion of all expression-related evidences, only three GO evidence codes IDA, HDA and IPI that represent direct experimental assays were considered. For the GO analyses of public microarray data the R package clusterProfiler(*52*) was used with default settings. The cutoffs used to define an enriched GO term include an adjusted P-value smaller than 0.01 and a fold enrichment greater than 2. To improve specificity only GO terms containing less than 200 genes were considered. The fold enrichment was calculated as (number of changed genes in the GO term / number of all changed genes) / (number of genes in the GO term / number of genes in all GO terms of the class). The GO semantic similarity scores were calculated by R package GOSemSim(*53*).

### Heritability analysis

Following a previous study(*54*), for each of the 195 genes the expression is expressed as *y* = *μ*1_*N*_ + *u* + *e*, where *y* is a vector of the expression level (log_2_RPKM) in the 24 segregants, *μ* is the mean expression level in the 24 segregants, 1_N_ is a vector of N ones, *u* is a vector of random additive genetic effects from the HAP4 locus, and *e* is a vector of residuals. The variance structure of an expression trait is written as 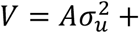 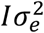, where *A* is relatedness matrix based exclusively on the HAP4 locus (1 for wild-type allele and 0 for null allele), *I* is identity matrix, 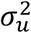 is additive genetic variance explained by HAP4 locus, and 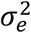 is error variance. Then, the value of *h*^2^_HAP4_ is equal to 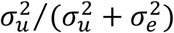. R package rrBLUP was used to estimate the variance components.

To test the statistical significant of an *h*^2^_HAP4_, 24 segregants were divided into two groups: 12 with the wild-type allele of HAP4 and 12 with the null allele of HAP4. We compared the expression levels of the focal gene between the two segregant groups using DESeq2. The obtained 195 raw P-values were adjusted for multiple testing using the Benjamini-Hochberg controlling procedure. An adjusted P-value smaller than 0.05 was considered significant.

### Analyze protein complexes and metabolic pathways

The public microarray data of ∼1,400 yeast gene deletion lines were obtained from a previous study(*26*), and P-values and fold changes (FC) provided in the data were directly used. Specifically, P < 0.05 and absolute FC > 1.2 were used to define genes with significant expression changes; if the number of significantly changed genes was over 1,500, a more stringent cutoff P < 0.01 was used. To avoid the effects of genes with ubiquitous expression responses we excluded from further analyses the top 10% genes that each show significant changes in at least 12% of the gene deletion lines. GO analyses were performed by R package clusterProfiler to define clustered and distributed effects for each deletion line, with the results summarized in Table S7. To examine the overlapped clustered effects between genes of the same protein complex or pathway, we only considered the deletion lines with at least 20 clustered effects, resulting in a set of 422 deletion lines suitable for further analyses.

Information of 518 protein complexes was obtained from a previous study (*35*). The KEGG pathways of the yeast *S. cerevisiae* were downloaded from KEGG website (https://www.genome.jp/kegg-bin/get_htext?sce00001). There are 54 complexes and 41 pathways each with at least two member genes found in the above defined mutant set.

For each protein complex or pathway, the overlaps of clustered effects and the overlaps of distributed effects were compared in number. The numbers of clustered effects and distributed effects of the involved genes were normalized to make the overlaps between the two effect types comparable. To estimate the confidence interval of a comparison we used random samplings. If clustered effects are less than distributed effects in all genes, which is true in most of the cases examined, we sampled (without replacements) a random subset of distributed effects to ensure the two effect types of a gene equal in number. If clustered effects are more than distributed effects in all genes, we sampled (without replacements) a random subset of distributed effects to ensure the two effect types of a gene equal in number. If the above consistent patterns do not exist, we sampled consistently from one side (either clustered effects or distributed effects) but with replacements for the gene with an insufficient number of effects on this side. For each complex or pathway 1,000 such random samplings were carried out to derive the 99% confidence interval, and an observed difference is called significant if it is not within the interval. Table S4 and Table S5 have details about the protein complexes and KEGG pathways examined, respectively.

### Analyze cell morphological traits

Diploid yeast cells were examined by following the protocol of previous studies with slight modifications(*38*, *39*). In brief, a single yeast colony was picked and cultured in YPD liquid overnight with shaking to the saturation phase. Then, 1.5μl culture was transferred to 100μl fresh YPD in a 96-well plate and grew for 3-4 hours at 30°C for SC strains or 25°C for SP strains. Cells were fixed with 3.7% formaldehyde solution. Cell wall was stained by FITC-ConA (fluorescein isothiocyanate-conjugated, concanavalin A, Sigma-Aldrich C7642). Cell nucleus was stained by hochest-mix (Thermo Fisher, Hoechst 33342 Solution) instead of DAPI to enhance the specificity. We did not stain actin because the dye Rhodamine phalloidin was not stable enough to support the following high-throughput automated image capturing which takes about 10 hours for scanning 96 wells of a plate. The stained cells were plated into a microplate (Greiner 781091) with ∼5.0×10^4^ cells per well and images were captured by IN Cell Analyzer 2200 (GE Healthcare) using the 60× objective lens.

Five SC lines (all diploid with BY4741 background: wild-type, *Δhap2*, *Δhap3*, *Δhap4* and *Δhap5*) and two SP lines (all diploid with N17 background: wild-type and *Δhap4*) were examined. Because the trait measuring is quite sensitive to batch effect, for each line we conducted 18-24 replicates of staining and image capturing. The images were analyzed by CalMorph(*38*, *39*) with default settings, and only 405 rather than 501 traits were extracted in this study because actin is not stained. At least 1,000 cells were captured and analyzed (with at least 100 informative cells for each cell-cycle stage) for a high-quality replicate. In the end, there were 13∼23 high-quality replicates for each of the lines included in further analysis. Trait values were compared between replicates of a gene deletion line and replicates of the corresponding wild-type line using T-test, and the resulting 405 P-values were adjusted for multiple testing using the Benjamini-Hochberg controlling procedure. Because of the many replicates included in the comparison, many traits showed a statistically significant but biologically negligible difference between wild-type and mutant lines. Hence, a trait is called affected by a gene only when the adjusted P < 0.05 and the difference between wild-type and mutant is large than 5%. Table S8 has complete information regarding the morphological trait analysis.

## Acknowledgments

We are grateful to financial support from NSFC (grant #31630042 and #91731302 to X. H.), technical support from Z. Zhou and X. Chen, and helpful discussions with C. Wu, J. Yang, J. Zhang, W. Qian, and Y. Zhang.

## Supporting Information

### Legends of supplementary tables

**Table S1:** RNA-seq-based gene expression levels for each HAP4 deletion and wild-type strain.

**Table S2:** RNA-seq-based expression changes (P-value and FC) of the 195 responsive genes in BY4741(*Δhap4*) and other deletion lines, with the values of *h*^2^_HAP4_ also included.

**Table S3:** RNA-seq-based gene expression changes after deleting HAP2, HAP3, HAP4 and HAP5, respectively, in BY4741.

**Table S4:** Summary of the analyses of protein complexes in this study.

**Table S5:** Summary of the analyses of KEGG pathways in this study.

**Table S6:** The affected morphological traits in a variety of gene deletion lines.

**Table S7:** Summary of the clustered effects and distributed effects defined in each mutant that has public microarray data.

**Table S8:** Summary of the trait information of each diploid gene deletion or wild-type yeast strain, with the 405 trait values, the number of examined cells, and the number of replications included.

**Fig. S1.**
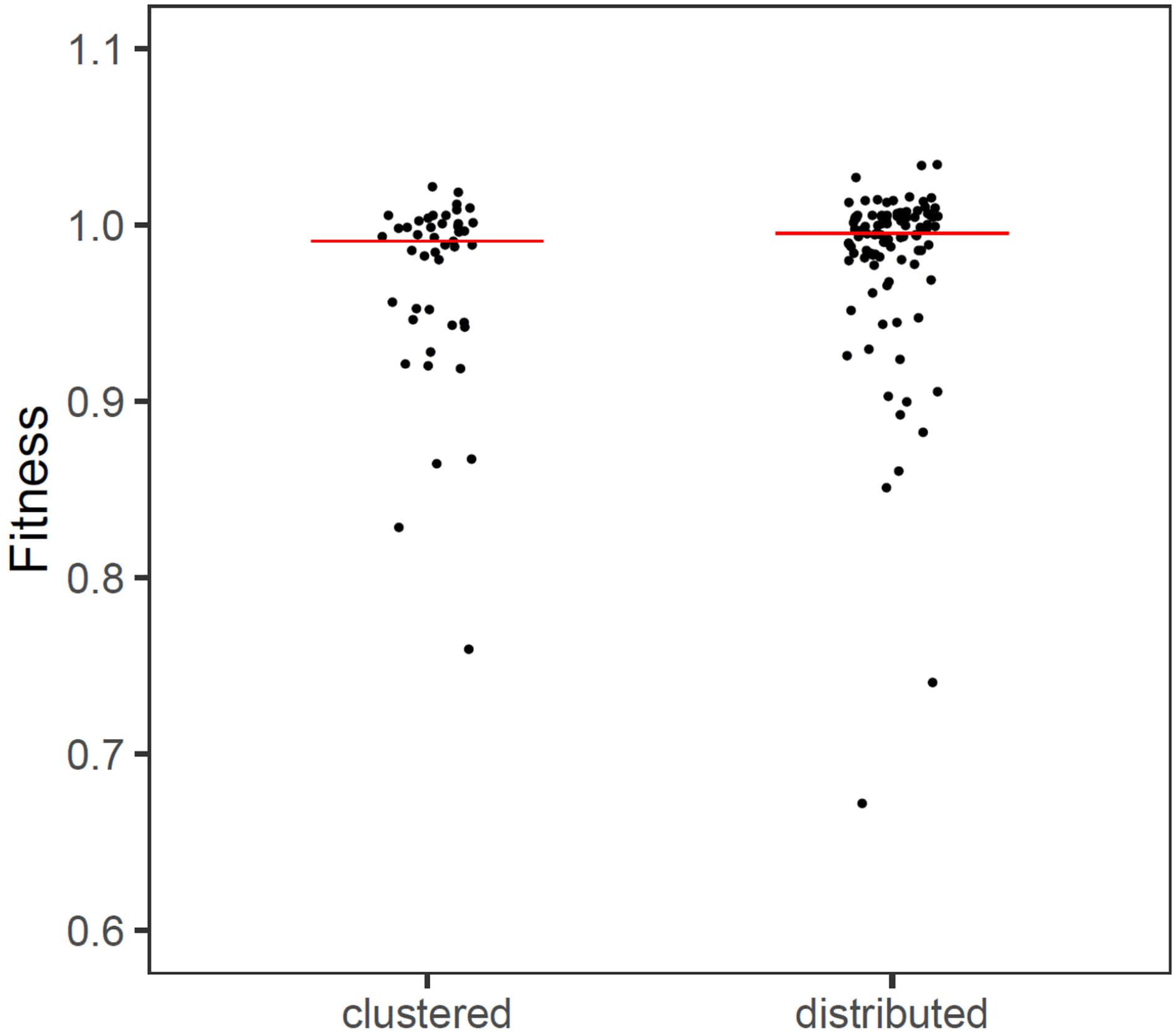
Fitness importance is comparable between the clustered responsive genes and distributed responsive genes defined in BY4741(Δhap4) (*P* = 0.10, Mann-Whitney U-test). Fitness importance of a gene is measured by the relative growth rate of the gene deletion line to wild-type. The horizontal line shows the median.

**Fig. S2.**
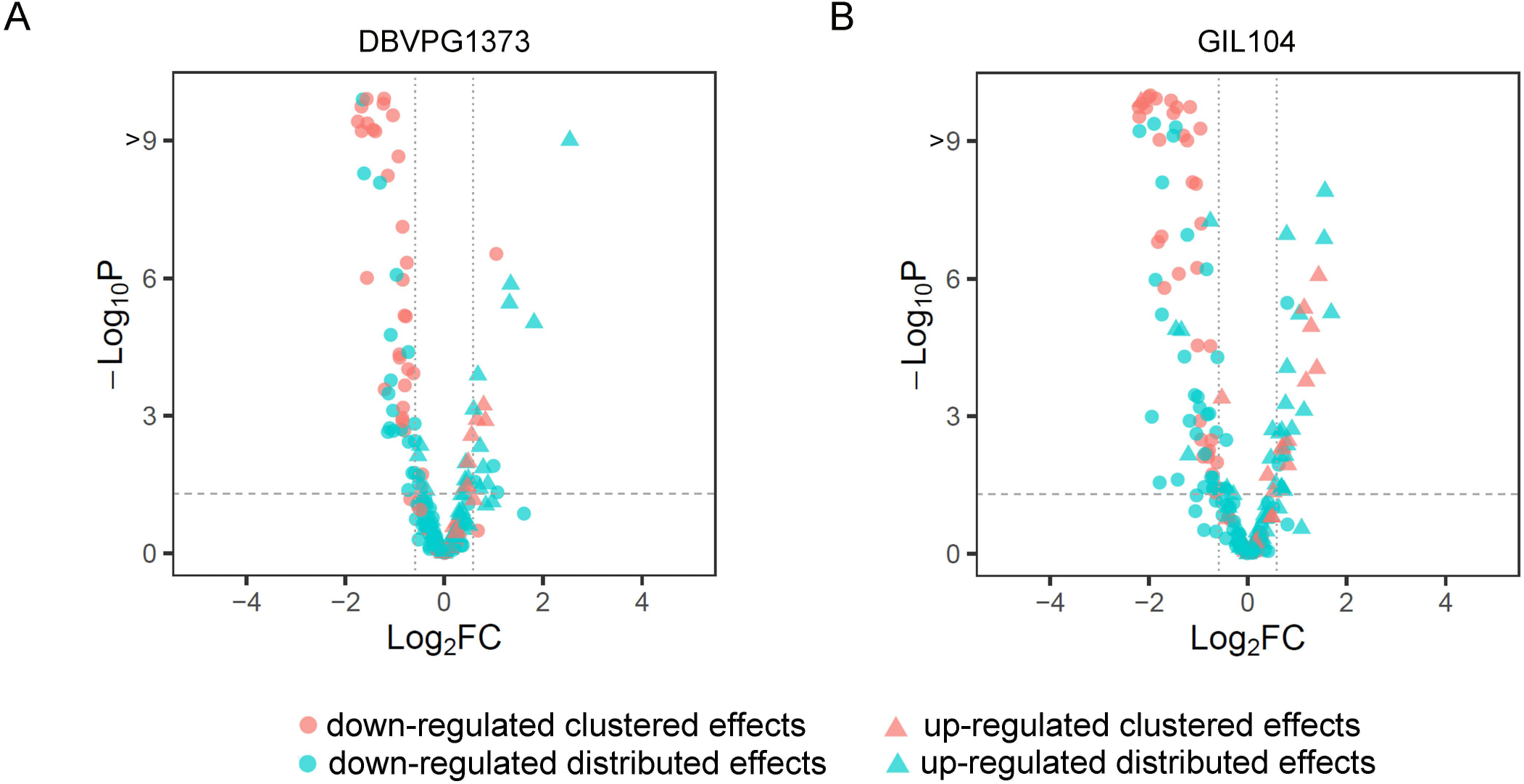
The intra-species conservation analysis of the HAP4 deletion effects. The 195 responsive genes defined in BY4741 (Δhap4) are examined with respect to their expression responses in *S. cerevisiae* DBVPG1373(Δhap4) and GIL104(Δhap4), respectively. The horizontal dashed line shows adjusted P = 0.05 and vertical dashed lines show log2FC = ±0.58. (cyan: clustered effects; red: distributed effects; cycle: down-regulated in BY4741(Δhap4); triangle: up-regulated in BY4741(Δhap4)).

**Fig. S3.**
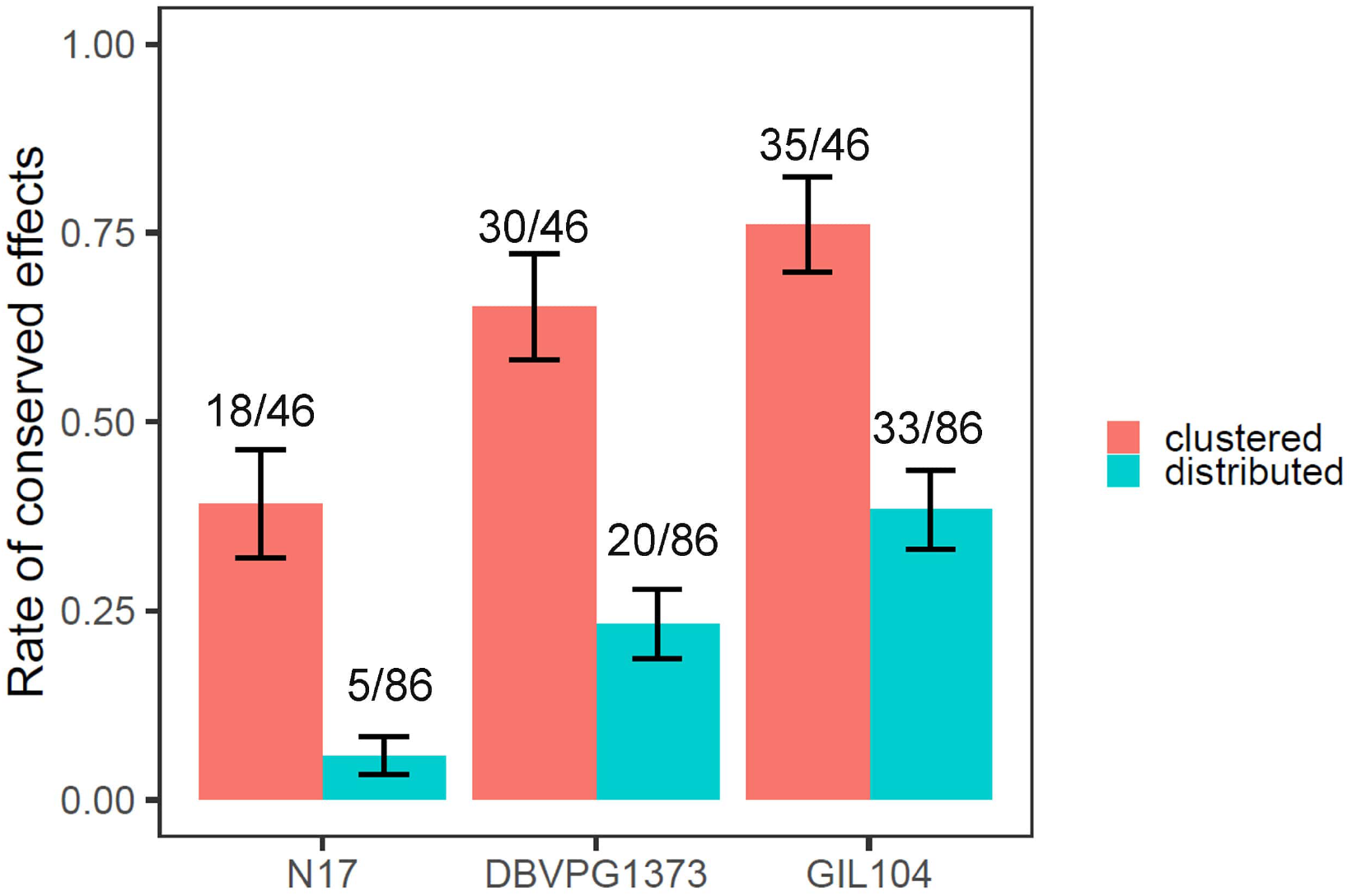
Conservation analysis of the HAP4 deletion effects by considering only genes with a strong expression level in wild-type BY4741. This analysis is to address the concern that lowly expressed genes in wild-type tend not to have detectable down-regulation due to technical bias. Hence, for the 195 responsive genes defined in BY4741 (Δhap4) only those with log2RPKM > 5 in wild-type BY4741 are considered here, leaving 46 clustered effects and 86 distributed effects. Error bars represent SE.

**Fig. S4.**
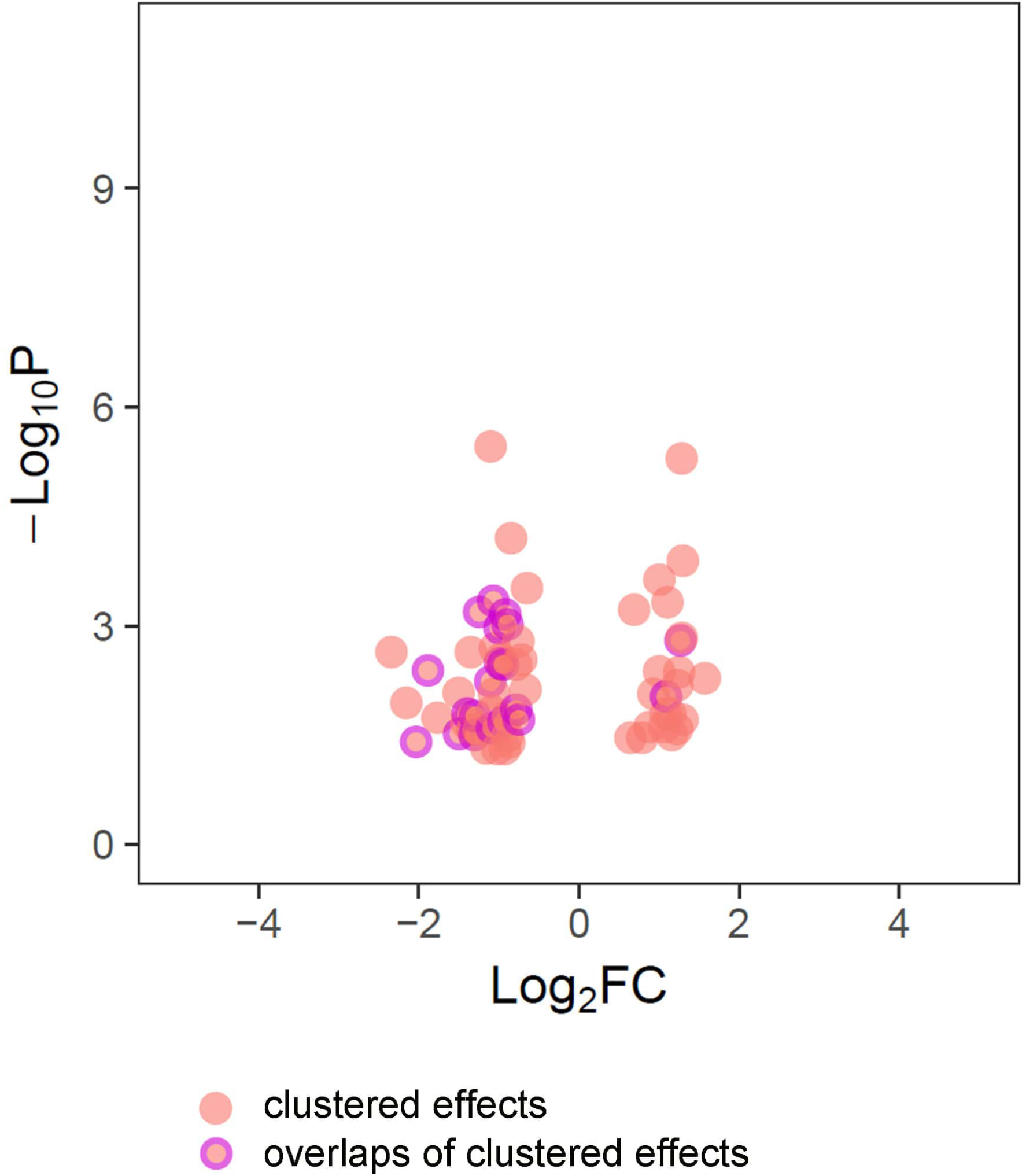
The 20 overlapped clustered effects are comparable to the rest 45 (65-20) clustered effects defined in BY4741 (Δhap4) with regard to their effect size. The differences are not statistically significant for both the P-values and fold changes observed in BY4741(Δhap4) (P = 0.75 and 0.51, respectively, Mann-Whitney U-test)

**Fig. S5.**
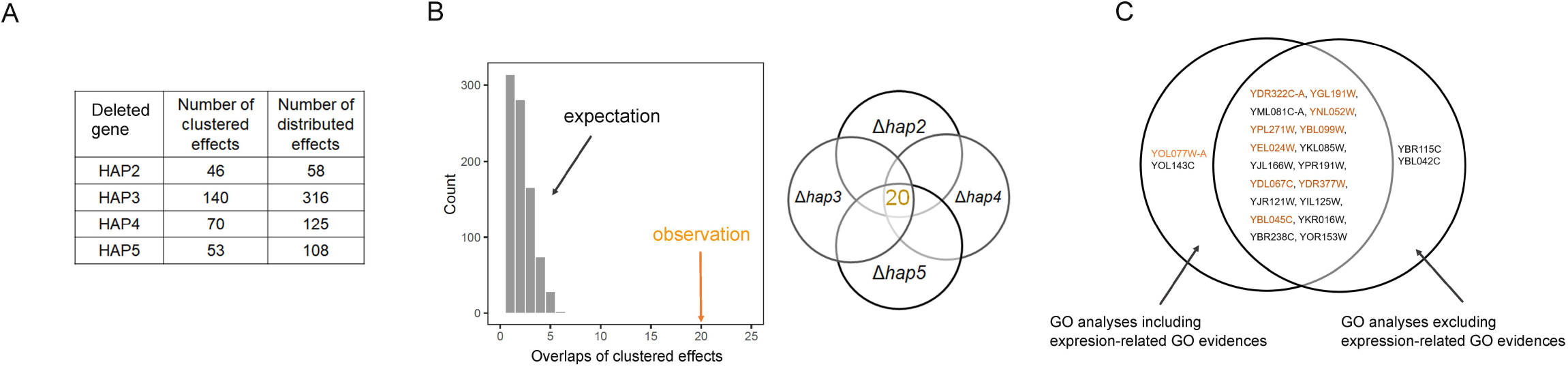
The strong overlaps of clustered effects of the HAP2/3/4/5 complex genes are not biased by expression-related GO evidences for annotating the deletion effects. (A) Only the evidences IDA, HOA and IPI are used to re-define the dustered and distributed deletion effects of the four genes, respectively. (B) There are 20 overlapped clustered effects for the four genes encoding the HAP2/3/4/5 tetramer, which is significantly higher than expectation. The expectation is estimated by random sampling of the distributed effects of the four genes to calculate overlaps, and 1,000 such simulations were conducted. (C) The overlapped clustered effects are largely the same before and after excluding expression-related GO evidences. Genes that are the direct target of HAP4 are highlighted in yellow.

**Fig. S6.**
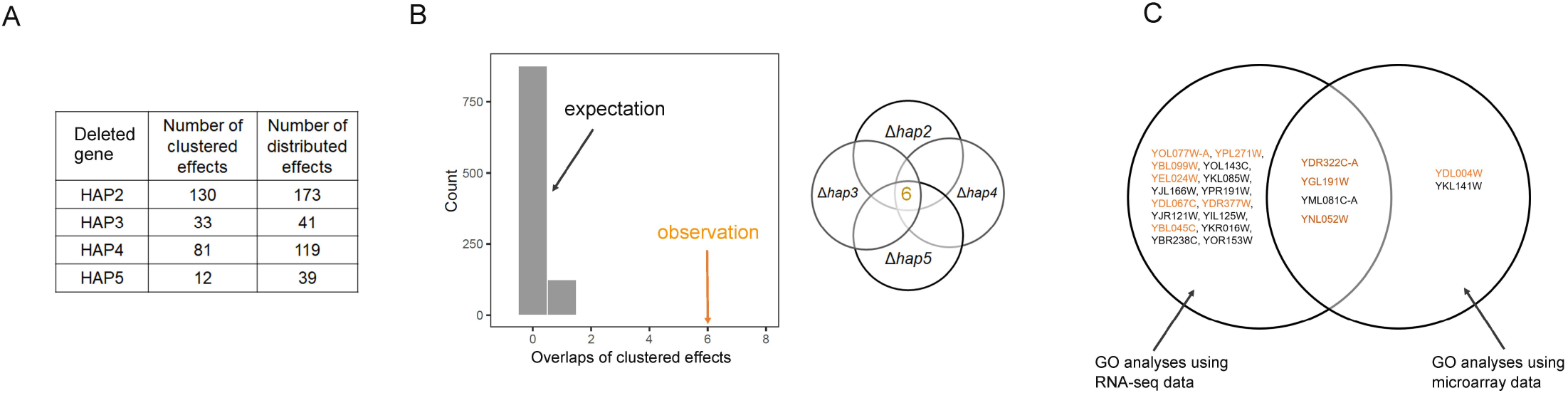
The enrichment of overlapped clustered effects in HAP2/3/4/5 complex is reproduced by using public microarray data of the four gene deletion lines. (A) Microarray-based expression data are used to re-define the clustered and distributed deletion effects of the four genes, respectively. (B) There are only six overlapped clustered effects for the four genes encoding the HAP2/3/4/5 tetramer, which is significantly higher than expectation. The reduced number is primarily due to the small number (12) of clustered effects observed in HAPS deletion. The expectation is estimated by random sampling of the distributed effects of the four genes to calculate overlaps; and 1,000 such simulations were conducted (C) Comparison of the six overlapped clustered effects defined using microarray data with the 20 overlapped clustered effects defined using RNA-seq data. Genes that are the direct target of HAP4 are highlighted in yellow.

**Fig. S7.**
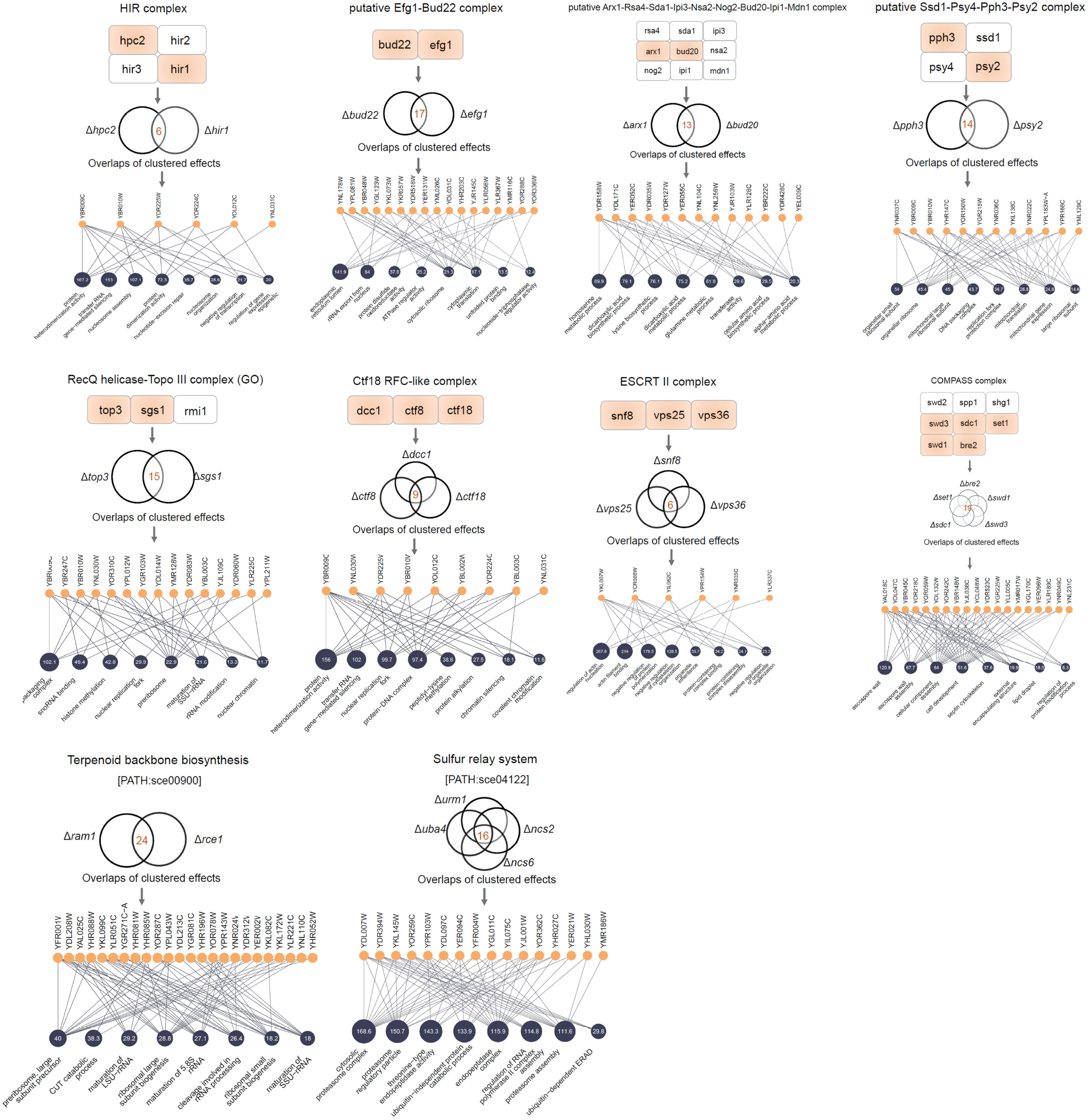
Clustered effects in general support related biochemistry understandings much better than distributed effects. Ten protein complexes or KEGG pathways each with a moderate number (6-24) of overlapped clustered effects are shown. The enrichment rate of overlapped clustered effects relative to distributed effects for each of the eight protein complexes is 9.0, 5.7, 3.4, 10.4, 3.8, 29.4, 26.4, and over 100, respectively. For the two KEGG pathways the enrichment rate is 15.4 (PATH:sce00900) and 551.7 (PATH:sce04122), respectively. The member genes of each complex or pathway with suitable expression data for the analysis are highlighted in orange. The orange circle each represents an overlapped clustered *effect*, and the blue circles represent the enriched GO terms of the overlapped clustered effects with the number inside showing the fold enrichment in the given term.

**Fig. S8.**
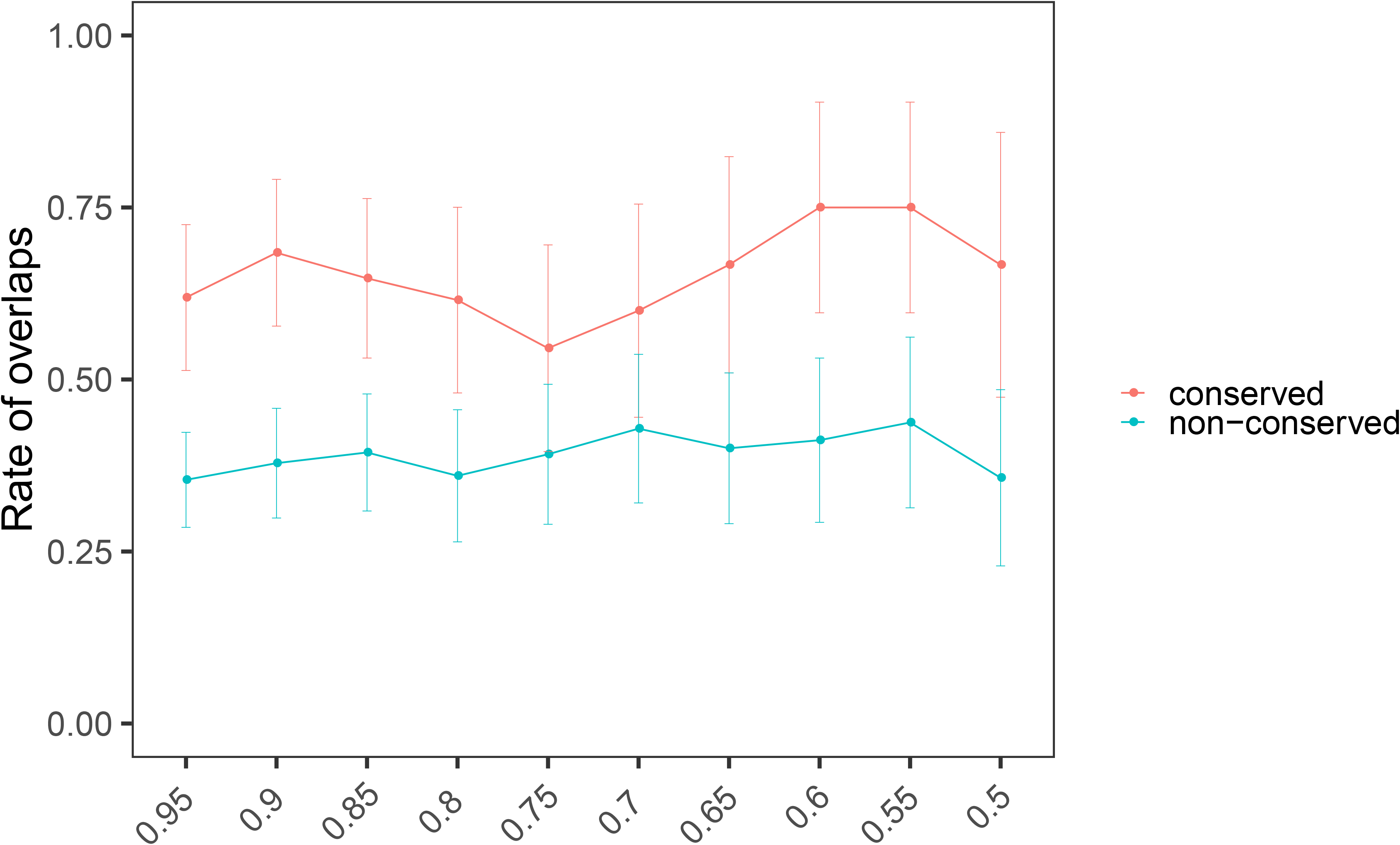
The estimated rate of overlaps cannot be explained by correlated traits. The Pearson’s R of all trait pairs is calculated using the trait values generated in ref. 4 for 4,718 yeast mutants. We then removed traits one by one from those with the highest absolute R until no two traits have R^2 greater than a threshold, which is set to be 0.95, 0.9, 0.85, 0.8, 0.75, 0.7, 0.65, 0.6, 0.55, and 0.5, respectively. The number of remaining traits are 346, 312, 277, 247, 223, 204, 185, 161, 153, and 127, respectively. Error bars represent SE.

**Fig. S9.**
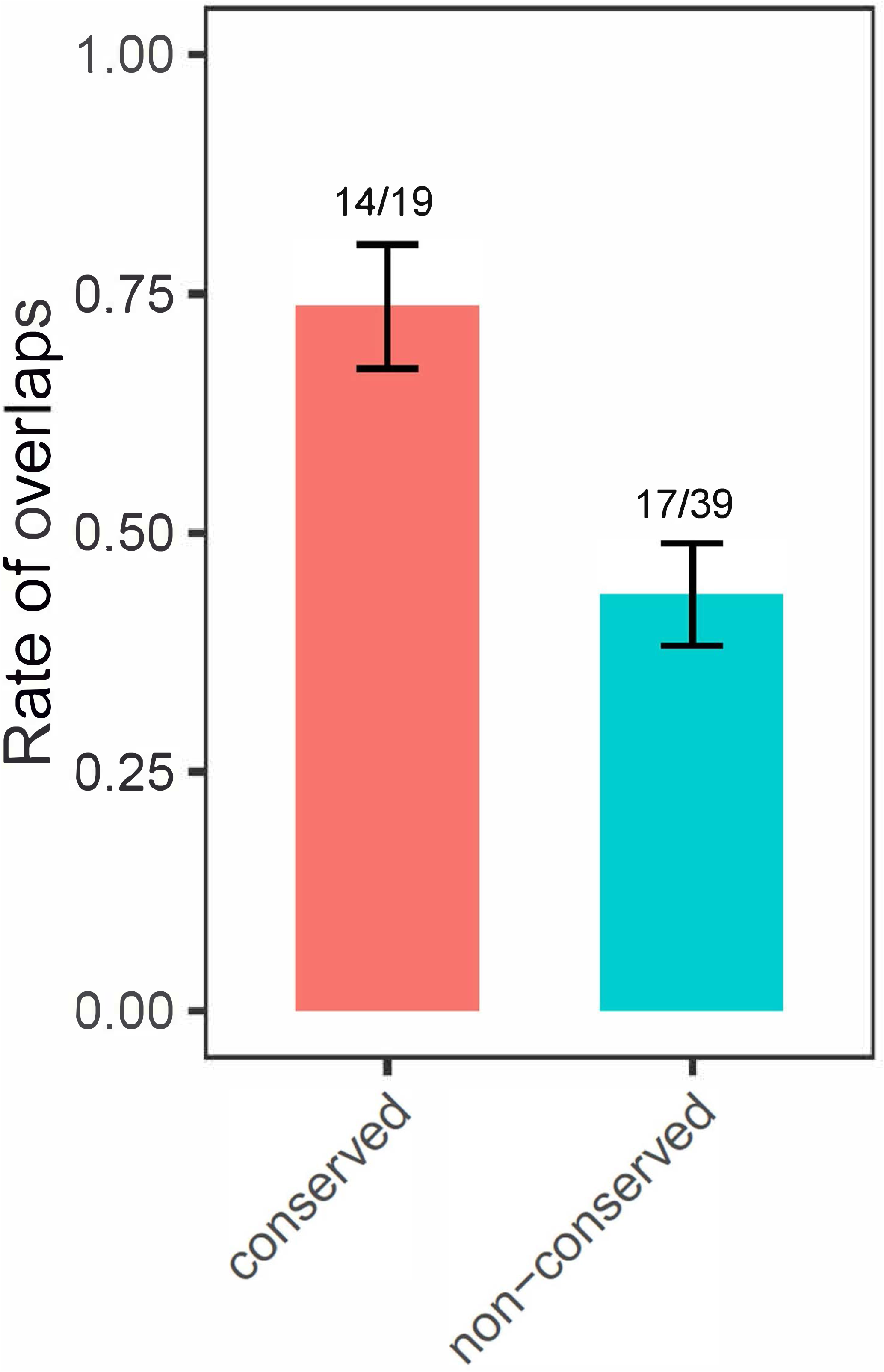
The comparison in Fig. 4B is robust against trait measuring noise. To address the potential technical bias that traits with large measuring noise tend to be both non-conserved and non-overlapping, only traits with measuring CV < 0.1 across the replicates in wild-type BY4741 are considered. This results in 58 traits that are significantly affected by HAP4 deletion in S. cerevisiae, among which 19 are conserved effects and 39 non-conserved effects. The rate of overlaps in the conserved set remains significantly higher than the non-conserved set (P = 0.029, one-tailed Fisher’s exact test). Overlaps refer to traits significantly affected by all four gene deletions in *S. cerevisiae.*

